# Asarinin Inhibits RANKL-Induced Osteoclast Differentiation by Targeting the p38/ERK–c-Fos–NFATc1 Axis

**DOI:** 10.64898/2026.01.21.700569

**Authors:** Lifang Zhang, Chengxu Xie, Xinyi Bao, Xiaohan Li, Heriberto Velez, Vishwa Deepak

## Abstract

Excessive osteoclast formation is a key contributor to pathological bone loss in disorders such as osteoporosis and rheumatoid arthritis. Asarinin, a natural lignan, has not previously been examined in the context of osteoclast differentiation. Here, we investigated the anti-osteoclastogenic effects of asarinin using RANKL-stimulated RAW264.7 cells. Asarinin significantly suppressed TRAP-positive multinucleated osteoclast formation under the tested conditions. Mechanistically, asarinin selectively inhibited RANKL-induced phosphorylation of p38 and ERK MAPKs, leading to reduced c-Fos expression and inhibition of NFATc1 nuclear translocation. In addition, asarinin disrupted actin ring formation in mature osteoclasts. Collectively, these findings identify asarinin as a pathway-selective inhibitor of osteoclast differentiation, targeting the p38/ERK–c-Fos–NFATc1 axis while sparing parallel signaling pathways.

## Introduction

Osteoclasts are multinucleated cells responsible for bone resorption, and their excessive activity leads to pathological bone loss in conditions such as osteoporosis, rheumatoid arthritis, and periodontal disease [1]. The receptor activator of nuclear factor-κB ligand (RANKL) is the primary cytokine that induces osteoclast differentiation by binding to its receptor RANK on osteoclast precursors [2]. RANKL-RANK interaction activates multiple downstream signaling cascades, including mitogen-activated protein kinases (MAPKs) such as p38, ERK, and JNK, as well as the NF-κB pathway [3]. These pathways converge to induce critical transcription factors including c-Fos and NFATc1. c-Fos, a component of the AP-1 complex, cooperates with NFATc1, the master transcription factor for osteoclast differentiation, to regulate osteoclast-specific gene expression [4].

Natural compounds with anti-osteoclastogenic properties have gained attention as potential alternatives to conventional bone resorption therapies due to their favorable safety profiles [5]. Importantly, pathway-selective modulators that target specific signaling cascades while preserving others may offer therapeutic advantages by minimizing compensatory mechanisms and off-target effects. Recent studies have identified novel molecular regulators of osteoclast differentiation, including intraflagellar transport proteins that modulate RANKL signaling [6]. Asarinin is a lignan isolated from *Asarum* species and has demonstrated anti-inflammatory and neuroprotective activities [7]. Notably, asarinin has been reported to modulate intracellular signaling pathways associated with inflammatory and stress responses, suggesting that it may influence differentiation programs regulated by MAPK-dependent transcriptional networks [8]. However, its effects on osteoclast differentiation and pathway selectivity have not been investigated. This study aimed to evaluate the anti-osteoclastogenic activity of asarinin and elucidate its pathway-specific molecular mechanisms using RANKL-induced RAW264.7 cells as an in vitro model.

## Materials and Methods

### Cell Culture and Reagents

Murine macrophage RAW264.7 cells were cultured in Dulbecco’s Modified Eagle Medium (DMEM; Gibco) supplemented with 10% fetal bovine serum and 1% penicillin–streptomycin at 37 °C in a humidified incubator with 5% CO_2_. Asarinin (purity ≥99%; CAS No. 133-04-0; HY-N0701) was purchased from MedChemExpress (NJ, USA). Recombinant mouse RANKL (462-TEC-010) was obtained from R&D Systems (MN, USA).

### TRAP Staining and Osteoclast Quantification

RAW264.7 cells were seeded in 96-well plates (5 × 10^3^ cells/well) and stimulated with RANKL (50 ng/mL) in the presence or absence of asarinin (10 μM) for 4-5 days. The culture medium containing RANKL and asarinin was refreshed every 2 days. Cell morphology and attachment were monitored microscopically throughout the differentiation period.

After differentiation, cells were fixed with 4% paraformaldehyde for 15 min and stained for tartrate-resistant acid phosphatase (TRAP) activity using a commercial TRAP staining kit (Servicebio, Hubei, China) according to the manufacturer’s protocol. TRAP-positive multinucleated cells (≥3 nuclei) were counted as mature osteoclasts under light microscopy (200× magnification). For each condition, at least five fields per well were analyzed and averaged, and osteoclast density was calculated by normalizing counts to the imaged area and expressed as osteoclasts per mm^2^. Osteoclast fusion index was calculated as the percentage of nuclei in multinucleated cells relative to total nuclei.

### Western Blot Analysis

Cells were lysed in RIPA buffer containing protease and phosphatase inhibitors. Equal amounts of protein (20-30 μg) were separated by SDS-PAGE and transferred to PVDF membranes. Membranes were blocked with 5% non-fat milk and incubated overnight at 4°C with primary antibodies against phospho-p38, total p38, phospho-ERK, total ERK, phospho-JNK, total JNK, phospho-p65, total p65, c-Fos, NFATc1, and GAPDH (Servicebio, Hubei, China). After washing, membranes were incubated with HRP-conjugated secondary antibodies and visualized using enhanced chemiluminescence (ECL). Band intensities were quantified using ImageJ software.

### Immunofluorescence Microscopy

For NFATc1 nuclear translocation analysis, cells cultured on glass coverslips were fixed with 4% paraformaldehyde, permeabilized with 0.1% Triton X-100, and blocked with 3% BSA. Cells were incubated with anti-NFATc1 antibody overnight at 4°C, followed by Alexa Fluor 594-conjugated secondary antibody. Nuclei were counterstained with DAPI. NFATc1 signals were visualized in red, while nuclei were counterstained with DAPI and pseudocolored in green. For actin ring visualization, cells were stained with Alexa Fluor 488-conjugated phalloidin. Images were captured using a Keyence microscope. All images were acquired using identical exposure settings within each experiment. The percentage of cells with nuclear NFATc1 and multinucleated osteoclasts with intact actin rings were quantified from at least five random fields. Data are expressed as mean ± standard deviation (SD) from at least three independent experiments.

### Statistical Analysis

Statistical significance was determined using Student’s t-test for two-group comparisons between RANKL and RANKL + asarinin groups. *P* < 0.05 was considered statistically significant.

## Results and Discussion

### Asarinin Suppresses RANKL-Induced Osteoclast Formation

To evaluate whether asarinin affects osteoclast differentiation, RAW264.7 cells were treated with RANKL in the presence of asarinin (10 μM). TRAP staining revealed that RANKL stimulation induced robust formation of TRAP-positive multinucleated osteoclasts in vehicle-treated controls (Figure 1). In contrast, asarinin treatment markedly reduced both the number of mature osteoclasts and the osteoclast fusion index (Figure 1). No overt changes in cell morphology or attachment were observed under the tested conditions. These results indicate that asarinin suppresses RANKL-induced osteoclast differentiation.

**Figure 1.**
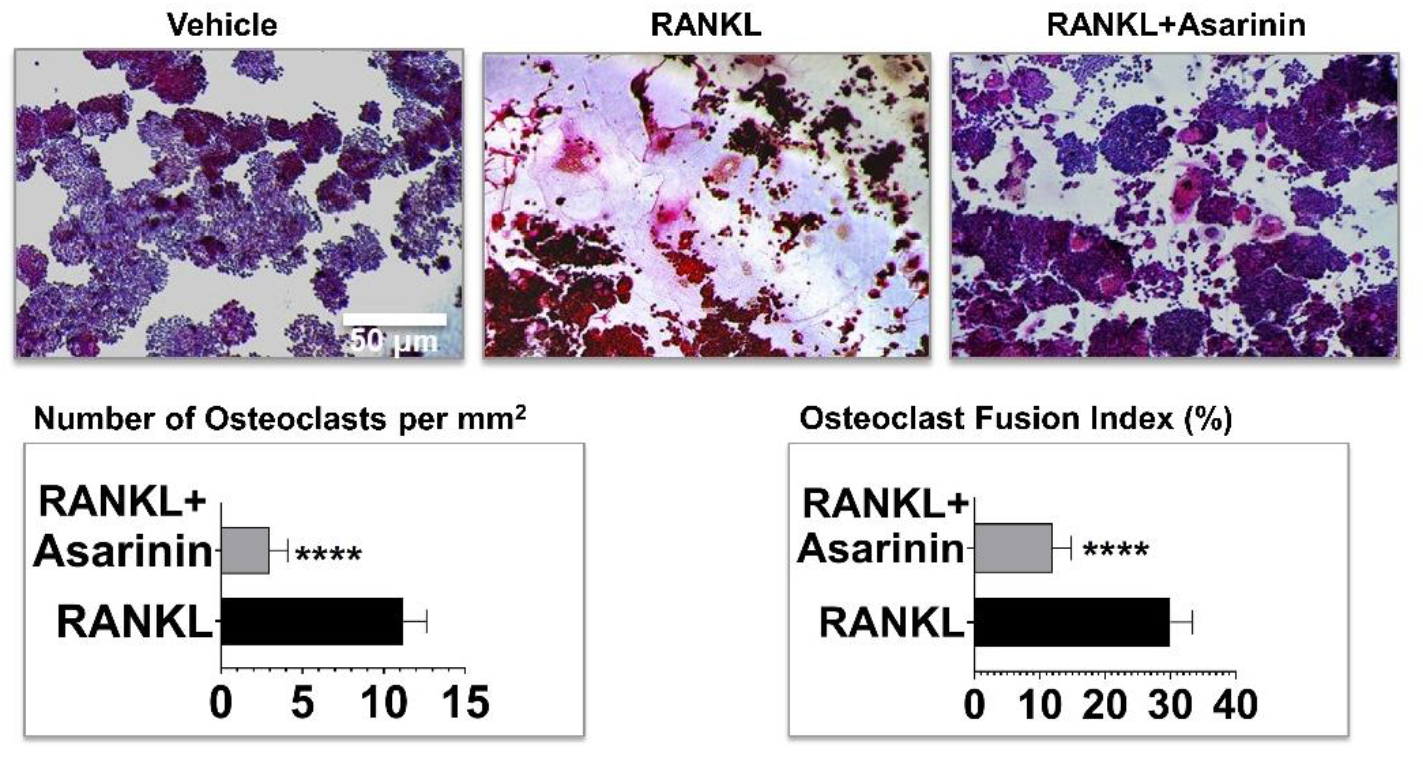
Asarinin suppresses osteoclast formation and fusion. RAW264.7 cells were cultured with RANKL (50 ng/mL) and treated with vehicle or asarinin (10 μM) for 5 days. Cells were fixed and stained for TRAP activity. Representative images show TRAP-positive multinucleated osteoclasts (purple). Osteoclasts were defined as TRAP^+^ cells containing ≥3 nuclei. Bar graphs show quantification of osteoclast number per mm^2^ and fusion index (%). Scale bar, 50 μm. Data represent mean ± SD from three independent experiments. \*\*\*\**P* < 0.0001 vs RANKL alone.

### Asarinin Exhibits Pathway Selectivity by Ameliorating p38/ERK while Preserving JNK and NF-κB Signaling

To understand the molecular mechanism underlying asarinin’s anti-osteoclastogenic activity, we examined its effects on RANKL-activated signaling pathways. Asarinin markedly suppressed RANKL-induced phosphorylation of p38 and ERK MAPKs (Figure 2A). In contrast, JNK phosphorylation and NF-κB activation (p65 phosphorylation) remained largely unaffected by asarinin treatment (Figure 2A), demonstrating clear pathway-specific modulation. This selective downregulation of p38 and ERK, while preserving JNK and NF-κB pathways, is particularly noteworthy as it suggests that asarinin acts through a specific molecular mechanism rather than general kinase inhibition. Such pathway selectivity may be therapeutically advantageous, as complete blockade of all RANKL-activated pathways could trigger compensatory mechanisms or unwanted side effects.

**Figure 2.**
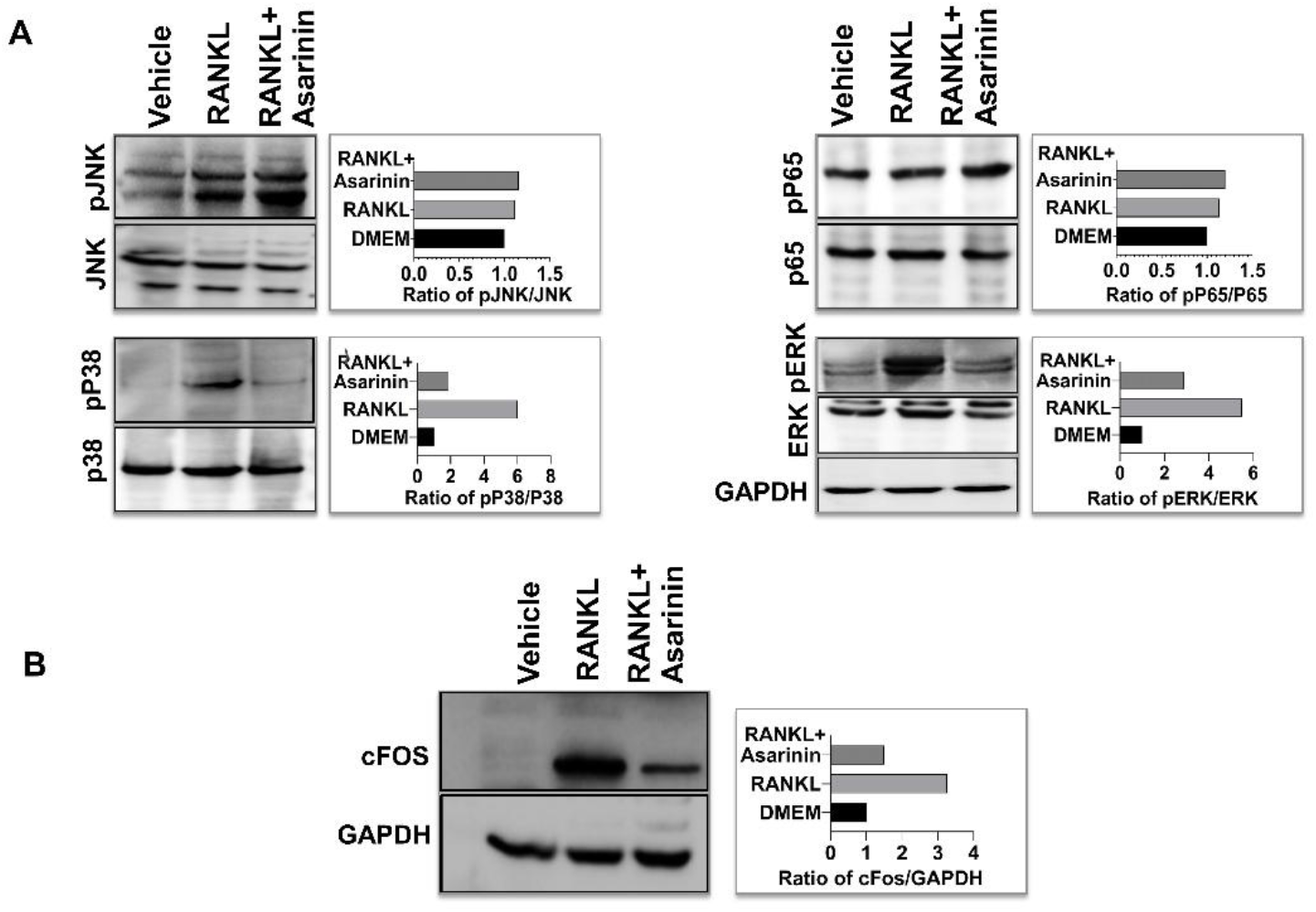
Asarinin selectively suppresses p38/ERK signaling, leading to reduced c-Fos expression. (A) RAW264.7 cells were treated with vehicle, RANKL (50 ng/mL), or RANKL plus asarinin (10 μM) for 10 min. Western blots show phosphorylation of p38 and ERK (reduced), whereas phosphorylation of JNK and p65 (NF-κB) was not significantly altered. Bar graphs show densitometric ratios normalized to total proteins. (B) RAW264.7 cells were stimulated with RANKL in the presence or absence of asarinin for 12 h. c-Fos expression was analyzed by Western blotting. GAPDH served as a loading control. Bar graphs show densitometric ratios normalized to GAPDH.

### Asarinin Suppresses c-Fos Expression and Prevents NFATc1 Nuclear Translocation

c-Fos is a key transcription factor induced predominantly by p38 and ERK MAPK signaling and is essential for NFATc1 induction during osteoclastogenesis [9]. Consistent with the selective inhibition of p38 and ERK, Western blot analysis revealed that asarinin significantly reduced c-Fos protein expression in RANKL-stimulated cells (Figure 2B), demonstrating the functional consequence of upstream MAPK modulation. NFATc1 is the master transcription factor that regulates osteoclast-specific genes, and its nuclear translocation is a prerequisite for osteoclast differentiation [10]. Immunofluorescence analysis revealed that RANKL stimulation induced robust nuclear accumulation of NFATc1 in control cells, with approximately 50% of cells showing nuclear NFATc1 localization (Figure 3A, lower panel). In contrast, asarinin treatment significantly inhibited NFATc1 nuclear translocation, with most NFATc1 protein remaining in the cytoplasm (Figure 3A, upper panel). This prevention of NFATc1 nuclear entry, coupled with c-Fos suppression, provides a comprehensive mechanistic explanation for asarinin’s potent anti-osteoclastogenic effect through the p38/ERK–c-Fos–NFATc1 axis.

**Figure 3.**
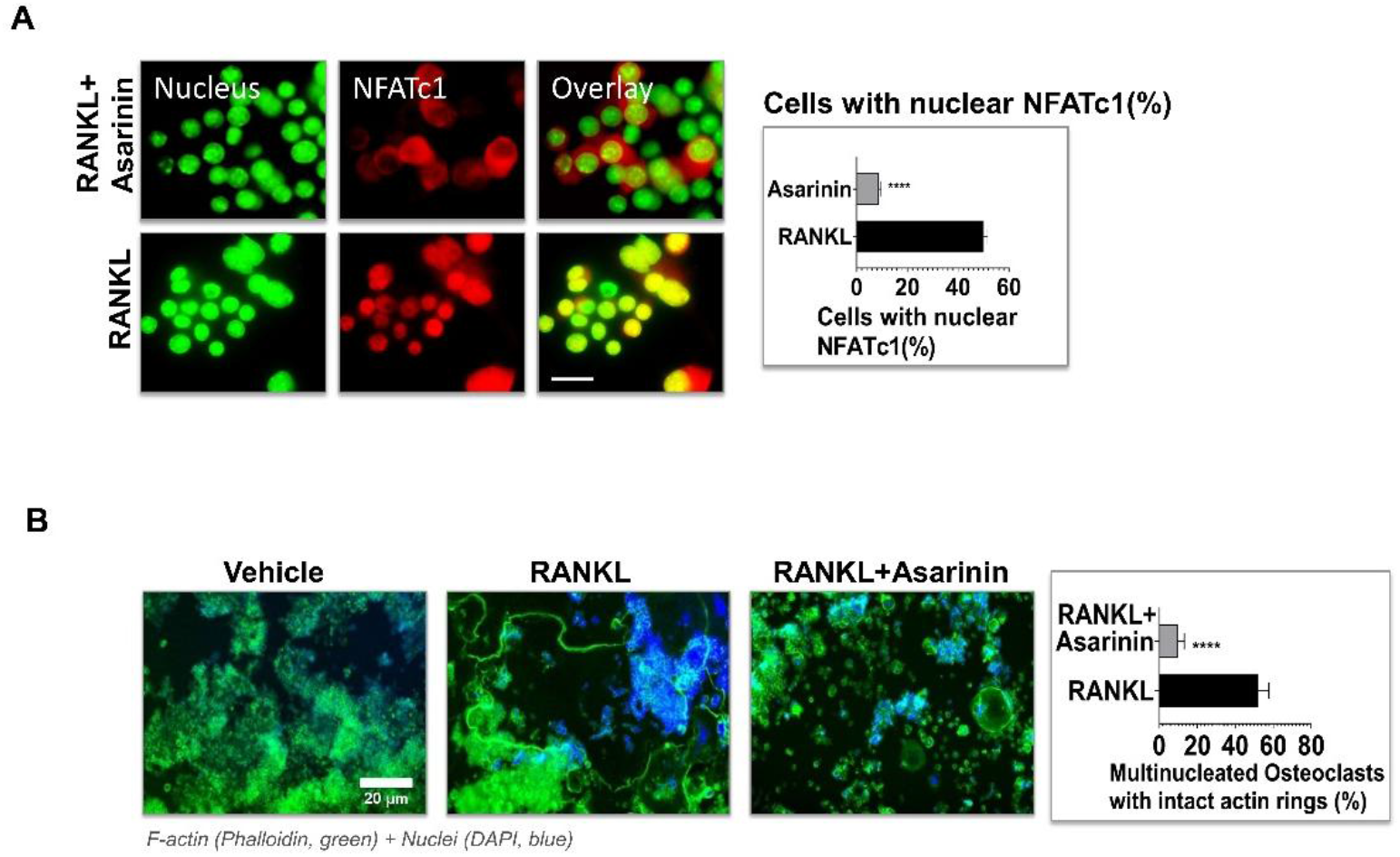
Asarinin prevents NFATc1 nuclear translocation and disrupts actin ring formation. (A) Immunofluorescence staining of NFATc1 (red) and nuclei (DAPI, pseudocolored green) in RAW264.7 cells treated with RANKL (50 ng/mL) in the presence or absence of asarinin (10 μM) for 3 days. Representative merged images show NFATc1 localization (yellow). Bar graph shows percentage of cells with nuclear NFATc1. Scale bar, 50 μm. (B) RAW264.7 cells were differentiated with RANKL in the presence or absence of asarinin (10 μM) for 5 days. F-actin (green) was visualized by phalloidin staining and nuclei (blue) were counterstained with DAPI. Representative images show actin ring structures in mature osteoclasts. Bar graph shows quantification of multinucleated osteoclasts with intact actin rings (%). Scale bar, 20 μm. Data represent mean ± SD from three independent experiments. \*\*\*\**P* < 0.0001 vs RANKL alone.

### Asarinin Disrupts Actin Ring Formation in Mature Osteoclasts

Functional osteoclasts require the formation of F-actin–rich sealing zones, or actin rings, to support bone resorption [11]. Phalloidin staining demonstrated prominent actin ring structures in RANKL-induced osteoclasts (Figure 3B). In contrast, asarinin treatment significantly reduced the proportion of multinucleated cells exhibiting intact actin rings (Figure 3B), indicating impaired osteoclast maturation and cytoskeletal organization. Disruption of actin ring formation further supports a functional inhibitory effect of asarinin on osteoclast development.

## Conclusion

In conclusion, this study demonstrates that asarinin is a pathway-selective inhibitor of RANKL-induced osteoclastogenesis, acting through multiple coordinated mechanisms: (1) selective suppression of p38 and ERK MAPK signaling while preserving JNK and NF-κB pathways, (2) reduction of c-Fos expression, (3) inhibition of NFATc1 nuclear translocation, and (4) disruption of actin ring formation. This pathway-selective profile distinguishes asarinin from non-selective kinase inhibitors and suggests a targeted mode of action that may confer therapeutic advantages by minimizing compensatory signaling and off-target effects. While conducted in vitro, these findings establish a mechanistic foundation for evaluating asarinin’s efficacy in animal models of osteoporosis and other osteoclast-mediated bone diseases.

## Author contributions

V.D. conceived and supervised the study and designed the experiments. L.Z. and V.D. performed the experiments. L.Z., C.X., B.X. and X.L. drafted the initial manuscript. H.V. provided technical support and methodological expertise. All authors reviewed and approved the final manuscript.

## Acknowledgments

We thank the core facilities of Wenzhou-Kean University for providing the necessary resources.

## Data availability

The authors confirm that all data generated during this study are included in this published article.

## Ethics approval and consent to participate

Not applicable.

## Consent for publication

Not applicable.

